# Understanding and Improving Word Embeddings through a Neuroscientific Lens

**DOI:** 10.1101/2020.09.18.304436

**Authors:** Sam Fereidooni, Viola Mocz, Dragomir Radev, Marvin Chun

## Abstract

Despite the success of models making use of word embeddings on many natural language tasks, these models often perform significantly worse than humans on several natural language understanding tasks. This difference in performance motivates us to ask: (1) if existing word vector representations have any basis in the brain’s representational structure for individual words, and (2) whether features from the brain can be used to improve word embedding model performance, defined as their correlation with human semantic judgements. To answer the first question, we compare the representational spaces of existing word embedding models with that of brain imaging data through representational similarity analysis. We answer the second question by using regression-based learning to constrain word vectors to the features of the brain imaging data, thereby determining if these modified word vectors exhibit increased performance over their unmodified counterparts. To collect semantic judgements as a measure of performance, we employed a novel multi-arrangement method. Our results show that there is variance in the representational space of the brain imaging data that remains uncaptured by word embedding models, and that brain imaging data can be used to increase their coherence with human performance.

## 1 Introduction

The most commonly-used approaches for representing semantic knowledge rely on corpus statistics. One such approach is a distributional semantic model (DSM), which uses word co-occurrence and frequency data to derive semantic representations (Landauer and Dumais, 1997). DSMs have become increasingly widespread since the introduction of word2vec by Mikolov et al. (2013) which learns vector representations for words based on their distributional properties. Existing word vector representations have been used to achieve impressive performance on many natural language tasks and they approach, if not surpass, human-level accuracy on several tasks such as sentiment analysis and named entity recognition. However, there still exists a significant gap between models making use of word embeddings and human performance on many other natural language understanding tasks (Nangia and Bowman, 2019). One approach to explaining the gap between humans and state-of-the-art NLP models is to assess the cognitive plausibility of the NLP models (Keller, 2010). This approach leads us to inquire whether existing word vector representations have any basis in how the human brain processes lexical semantics.

The seminal paper published by Mitchell et al. (2008) first demonstrated that word vectors sourced from corpus statistics could be used to predict participants’ functional magnetic resonance imaging (fMRI) responses when they were shown concrete nouns. Follow-up work has replicated the original paper’s results with other brain imaging modalities such as magnetoencephalography (Sudre et al., 2012). While the semantic features of the word vectors Mitchell et al. used were defined by cooccurrences with 25 manually-selected verbs, state-of-the-art DSMs have been used to improve the original results (Anderson et al., 2017; Abnar et al., 2018). More recently, brain imaging data during the onset of a word in the context of a sentence (or a narrative) have been linked to decoding models employing word embeddings (Wehbe et al., 2014; Anderson et al., 2016a; Pereira et al., 2018). Collectively, these findings cohere with evidence which suggests that a continuous semantic space exists in the human brain (Huth et al., 2012; Huth et al., 2016).

In order to continue building on this existing body of work, we must address two important questions which have largely gone unanswered:

1. Do the representational structures of words derived from existing word embedding models differ from those in the brain?
2. If their representations differ, can we use features from the brain to improve the performance of existing models for creating word embeddings in NLP?

To establish a performance metric by which to answer the second question, we use a novel technique for acquiring semantic judgements of textual stimuli: the multi-arrangement method (Kriegeskorte and Mur, 2012). The multi-arrangement method asks participants to directly model the representational space we are interested in, while overcoming many of the issues associated with the existing methods for collecting semantic judgements in NLP.

Following this introduction, we first describe the brain imaging data and the word vector representations to be used in this study (Sections 2 and 3). Then we describe how we compare the representational structure of individual words for the brain imaging data and the word vector representations, using a method known as representational similarity analysis (Section 4). We subsequently introduce the motivation behind our use of the multi-arrangement method, and we describe the details of the behavioral data we collected (Section 5). Using these data as an evaluation metric, we then compare the performance of the word vectors constrained to the features of the brain imaging data with the performance of the unmodified word vectors (Section 6). We ultimately find variation in the brain imaging data (beyond noise) that remains uncaptured by the word embeddings, and that brain features can be used to improve the performance of existing word embedding models (to be discussed in Section 7).

## 2 Brain Imaging Data

In our paper, we use the previously collected fMRI dataset from Mitchell et al. (2008)^1^. The fMRI data were collected from 9 participants as they were asked to name properties for each word shown. Each fMRI image was captured while a participant was shown a concrete noun and its corresponding line drawing. Concrete nouns belonged to one of 12 semantic classes (e.g. mammals, buildings, tools) and there were 5 such exemplars presented for each semantic class. Each concrete noun was displayed six times, in random order, resulting in 360 fMRI images per participant. The fMRI data we use for analysis were the same as in the original paper except for minor differences in preprocessing described below.

### 2.1 Preprocessing

When evaluating their encoding model, Mitchell et al. (2008) used only the fMRI data from the 500 voxels^2^ from the whole brain with the most stable response profiles for the 60 concrete nouns, across all six presentations. Mitchell et al. approximated the stability of each voxel as the average Pearson correlation between its responses across each possible pair of the six runs. This stability-based approach to voxel selection is susceptible to including regions which respond identically to all stimuli (i.e. this approach would select for voxels whose activations do not vary across words). Instead, we use a reliability-based approach for voxel selection following the methodology introduced by Tarhan and Konkle (2019). We first assess the reliability of each voxel by computing the Pearson correlation of its response profile to all 60 words across even and odd runs. This results in one r-value (i.e., measure of reliability) for each voxel in a given participant’s fMRI data. We then consider a range of possible thresholds for the r-value (between 0 and 0.95). Across both even and odd runs, we calculate the reliability of the response patterns in the voxels included in the given threshold for each word. This calculation produces a series of values of r and their corresponding reliability across response patterns for each word. After analyzing all such series of values across all participants, we find that the threshold for *r* which optimized both the reliability and the coverage of the data was r = 0.25^3^. Since this threshold is selected without considering which words garner the greatest responses from the voxels (or independently of any specific relationship among the words in the dataset more generally), there is no need to perform voxel selection separately for each validation set^4^. Moreover, this value for r is similar to the optimal value of r = 0.30 that Tarhan and Konkle (2019) found for the fMRI datasets they evaluated in their paper.

We additionally transform each word’s response from each voxel into a z-score as was also performed by Anderson et al. (2016b) in their work with the data from Mitchell et al. (2008). This is done in order to remove amplitude differences between conditions and runs. Beyond this normalization procedure, we form the representation for each concrete noun by averaging its standardized response pattern over the selected voxels across all six presentations of the noun.

### 3 Word Vector Representations

When considering which word vector representations to use in this work, we decided against contextualized word embeddings for several reasons. We cannot make use of one of the main advantages of contextualized word embeddings—context—due to the nature of our fMRI data where an isolated noun is displayed during each trial. We could not clearly justify introducing context into our dataset (e.g. by providing the respective Wikipedia entries for each concrete noun) while maintaining a fair comparison with the remaining word vector representations. For these reasons, we have chosen to use word vector representations which are not contextualized. The most commonly-used such word vector representations are:

- **word2vec.** Word2vec uses a skip-gram model which learns to predict words using their context (Mikolov et al., 2013). We use the vectors trained on 100 billion words from the Google News dataset^5^.
- **GloVe.** GloVe is a global context method which makes use of word-word co-occurrence statistics across an entire corpus (Pennington et al., 2014). We use the vectors trained on 42 billion words from the Common Crawl dataset^6^.
- **fastText.** FastText is an extension of word2vec which treats each word as a group of character n-grams, thereby learning to account for morphological information (Bojanowski et al., 2017). We use the vectors trained on 16 billion words from the Wikipedia 2017, UMBC webbase, and the statmt.org news datasets^7^.

All the pre-trained word vectors we use are of length 300. Despite their qualitative differences, all three word vector representations rely on distributional semantics.

### 4 Comparing the Representational Spaces of the Brain and Word Vectors

In order to compare the representational spaces of the brain and word vectors, we will use representational similarity analysis (RSA).

#### 4.1 Representational Similarity Analysis

RSA was first introduced by Kriegeskorte et al. (2008) and has become a popular procedure in the field of cognitive neuroscience for relating stimuli representations from various modalities with brain imaging data. The procedure relies on the idea that correlating pairwise distance judgements across qualitatively distinct models (e.g. brain imaging data and computational implementations) can serve as a proxy for determining the extent to which the representational spaces of the models are aligned. In our paper, we will adopt the RSA methodology outlined in Nili et al. (2014), which introduced a toolbox implementing the RSA procedure.

Before we can perform our comparative analysis of the representational spaces of the participants’ fMRI data and the word vectors, we must compute the representational spaces themselves. For each of the concrete noun representations, we compute a representational dissimilarity matrix (RDM) wherein each entry denotes the pairwise distance (defined as 1–*ρ,* where *ρ* is the Pearson correlation) between the corresponding pair of nouns under the given representation. Thus, each representational space will be defined by the upper triangular region of a symmetric RDM with dimensions 60×60.

To determine how related a word vector’s RDM is to the participants’ brain imaging data RDMs, we calculate the Spearman correlation^8^ between the RDM of the word vector and a given participant’s RDM, and we average the results across all participants. We evaluate the significance of the correlations using a one-sided Wilcoxon signed-rank test (against the null hypothesis of a correlation of 0). Because the Wilcoxon signed-rank test is nonparametric, there is no assumption made regarding how normally-distributed our data is. The results of these tests are shown in Figure 1.

**Figure 1:**
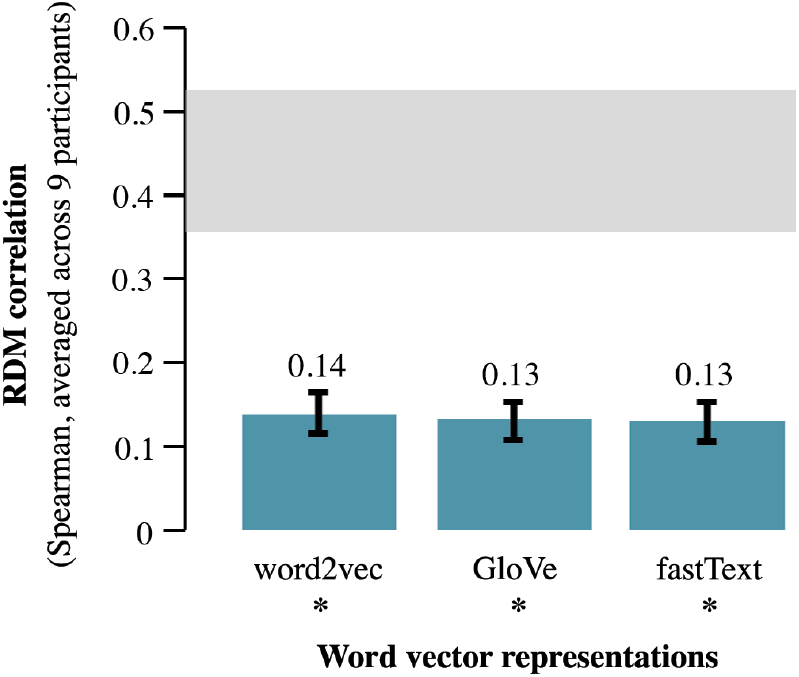
Results of the comparative analysis of RSA. The star below each word vector representation denotes if it is significantly related to the participants’ fMRI data, *p* < 0.05. Here, the error bars above each word vector representation indicate the standard error of the mean resulting from the variation across participants. The values above the word vector representations specify the average correlation between the participants’ RDMs and the respective word vector representation. The shaded area indicates the lower and upper bound of the noise ceiling (i.e. the performance we would expect from a true model of the underlying representational space, considering the noise present in the data). The upper bound is estimated using the correlation between each participant’s RDM and the RDM averaged across all participants (including the participant we are correlating with), while the lower bound is computed by correlating each participant’s RDM with the RDM averaged across all remaining participants^9^. The visualization has been adapted from the Matlab toolbox^10^ provided by Nili et al. (2014).

Each word vector representation’s RDM has a correlation significantly greater than 0 (p’s **=** 0.002). Thus, all three word vector representations’ RDMs are significantly related to those of the participants’ fMRI data. However, the difference between the correlation of the word vector representations and the correlation we would expect from a true model (accounting for noise) suggests that much of the representational space of the brain imaging data is not aligned with that of the word vector representations.

We then assess if the word vector representations differ in their relatedness to the participants’ fMRI data. For each pair of word vector RDMs, we perform a two-sided Wilcoxon signed-rank test against a null hypothesis that both RDMs’ correlations with each of the participants’ RDMs are equal (due to the number of samples provided, the two-sided Wilcoxon signed-rank test is an exact test). After correcting the resulting significance values for multiple tests using the Benjamini-Hochberg method (Benjamini and Hochberg, 1995), there is no significant difference in the degree to which each of the word vector representations is related to the brain imaging data (p’s > 0.05). This result may be explained by our understanding that all the word vectors compared are DSMs.

Returning to our first question, we can conclude that the representations of the word embeddings are significantly related to the representations of the brain imaging data. However, there is still variation beyond noise in the brain imaging data that these methods do not capture.

### 5 Introducing a Novel Semantic Judgement Acquisition Technique

Before we can evaluate whether features from the brain can be used to improve the performance of existing word embeddings, we must first define a metric by which we will judge performance (Pereira et al., 2016). A common approach to evaluating vector space representations in NLP is to compare the semantic relations predicted by the distances between word vectors with human-rated relatedness/similarity scores. However, the largest publicly-available word similarity/relatedness benchmark has minimal overlap with the word pairs in our dataset (it only contains scores for 13 of the 1770 possible word pairs). There are also several shortcomings to the existing methods of collecting participant similarity and relatedness judgements for textual stimuli. Problems include conflating between types of relations (e.g. synonymy and antonymy) in the case of word association norms, relying on heuristics (e.g. “good-bad”, “weak-strong”) in the case of the semantic differential technique, and being affected by bias from both word order and context in the case of pairwise similarity judgements.

#### 5.1 The Multi-arrangement Method

We make use of a new approach for collecting participants’ semantic judgements in NLP referred to as the multi-arrangement method (Kriegeskorte and Mur, 2012). The multi-arrangement method requires participants to perform arrangements of several subsets of stimuli. Each subset is designed to collect an equal amount of evidence for the dissimilarity between each pair of stimuli. A dissimilarity matrix for the stimuli is then computed from the redundant distance information in the multiple arrangements. A demonstration of the multi-arrangement method is shown in Figure 2. In past work, the multi-arrangement method has been primarily used to collect semantic judgements of images and visual object categories (Mur et al., 2013; Dobs et al., 2019; Karimpur et al., 2019). This approach avoids many of the issues present in existing methods for collecting human-rated similarity and relatedness scores through asking participants to directly model the representational space for the concepts we desire.

**Figure 2:**
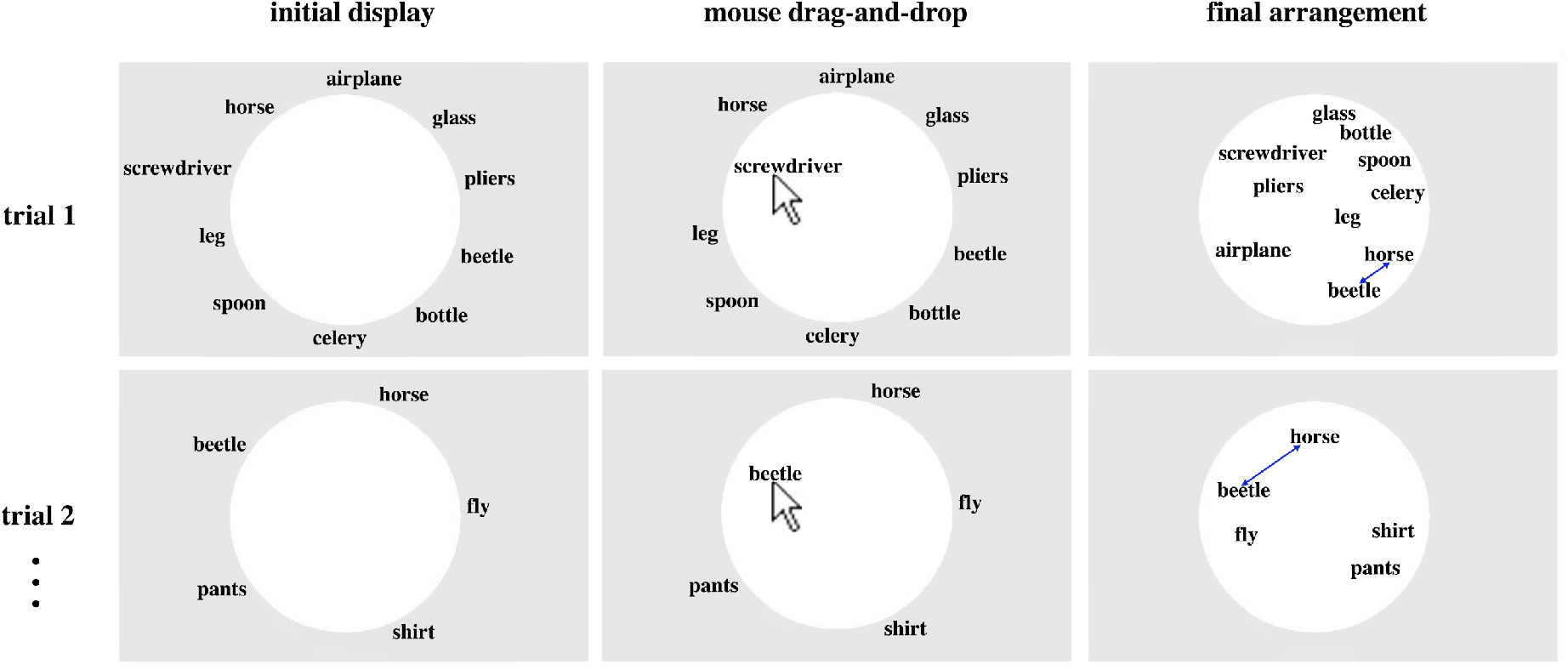
A demonstration of the multi-arrangement method using a toy example is shown. A participant is first presented with an initial set of stimuli to arrange (words in this case). Following a drag-and-drop scheme, the participant then arranges the stimuli based on their perceived relationship (i.e. more similar stimuli are arranged closer together, while more dissimilar stimuli are arranged further apart). The subsequent subset of stimuli to arrange is determined by the “lift-the-weakest” algorithm which aims to collect an equal amount of evidence of dissimilarity for each stimulus pair. A dissimilarity matrix is then computed using the redundant information present in the participant’s multiple arrangements. This figure has been adapted from the one provided in Kriegeskorte and Mur (2012).

Several authors from Kriegeskorte and Mur (2012) and follow-up work with the method (Charest et al., 2014) created the Meadows Research^11^ platform, which implements the multi-arrangement method in an online tool. For our paper, we recruited participants in-person and collected semantic judgements using the multi-arrangement method (as implemented on the Meadows Research platform). To the best of our knowledge, this is the first time the multi-arrangement method has been used in NLP to collect semantic judgements from participants.

#### 5.2 Behavioral Data

Ten healthy individuals (3 identified as female and 7 as male; mean age: 21.4 years, standard deviation: 1.50) volunteered to participate in our experiment. All participants were fluent speakers of English. Each participant had 60 minutes to complete several arrangements (as many as they comfortably could) of the concrete nouns in our dataset. Following the multi-arrangement method, an RDM is computed for each participant based on the redundant information in their arrangements. We average the RDMs across all ten participants, providing us with 10 semantic judgements ^12^ for each of the 1770 word pairs in our dataset. This RDM of pairwise dissimilarity judgements will form our evaluation metric.

### 6 Improving Existing Word Vector Representations

Now that we have established a metric by which to evaluate word vectors, we can proceed with determining whether features from the brain can be used to improve the performance of existing word vector representations. We will answer this question by first mapping the word vectors onto the fMRI data using linear regression. We will evaluate this mapping under a cross-validation procedure to assess if the distances it predicts between previously unseen words correlate better with the behavioral data than existing word vector representations.

Following from the analysis of Mitchell et al. (2008), we decide to learn mappings using ridge regression, a variant of multiple linear regression that uses a least squares loss function and L2 regularization. For every word vector representation and each participant’s fMRI data, we learn a mapping between the features of the word vectors and the voxels in the participant’s preprocessed fMRI data. This mapping is used to predict the fMRI responses for previously unseen nouns. Given that no word vector representation entirely captures the representational structure of the fMRI data, the learned mappings may allow for the uncaptured representational structure in the fMRI data to supplement the word vectors in predicting the variance of the behavioral data. In order to evaluate the learned mappings, we use a leave-two-out cross validation procedure similar to that originally used by Mitchell et al. (2008).

For each combination of word vector representation and participant, we train a ridge regression model with the word vectors and the fMRI data for 58 of the 60 nouns, repeating this training process for every such combination of left-out words. We then use this regression model to predict the fMRI images of the two left-out words. Then we can compute the cosine distance between those two predicted fMRI images, and have that serve as the predicted relatedness score between the left-out words. After performing this operation for all combinations of left-out word pairs, 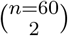, the resulting 1770 predicted relatedness scores can then be correlated with the behavioral data we collected earlier to assess the performance of the mapping between the given word vector representation and participant.

The results after evaluating the mapping learned between each word vector representation and each participant’s fMRI data are shown in Table 1^13^.

**Table 1:**
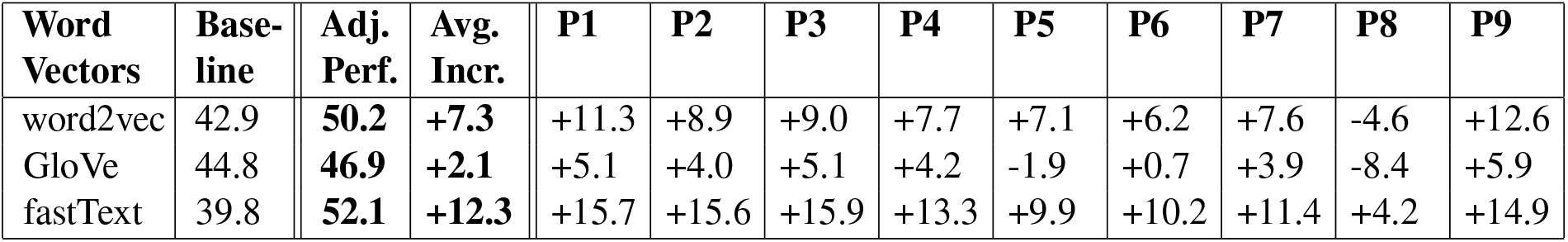
The performance of each word vector representation before and after ridge regression is shown. Immediately on the right of each word vector representation is the performance of the corresponding vectors before ridge regression (i.e. the Spearman correlation between the word vector representations themselves and the behavioral data we collected earlier). The following two columns display the performance of each word vector representation and its respective increase from the baseline after ridge regression, averaged across all participants. The rightmost columns show the change in performance when the given word vector representation is mapped onto the given participant’s fMRI data (the specific participant is denoted at the top). The values displayed are multiplied by 100 for readability.

We assess the significance of the ridge regression results by comparing it with those of an empirical distribution of null models (in a manner similar to that performed by Mitchell et al. (2008) to assess the significance of their regression results). We perform our permutation testing by first taking the fMRI data of one of our participants at random, and shuffling the 60 word labels. A model is then trained and tested (by the leave-two-out cross-validation procedure described earlier) using these data and the word vectors corresponding to the 60 word labels in the original (non-permuted) order. We train 1,000 such models for each word vector representation in order to form an empirical distribution. From this distribution, we find that each of the mappings we learned between the participant fMRI data and the word vector representations exhibit a correlation greater than 99.9% of null models in the empirical distribution. We therefore assign a significance value of *p* < 0.001 to each of our participant models for being more strongly correlated with the human semantic judgements than the 99.9^th^ percentile model in the null distribution for each word vector representation.

We further test the significance of the improvements across participants with respect to each word vector representation’s baseline performance using a one-sided Wilcoxon signed-rank test (against the null hypothesis of no improvement in performance)^14^. We find that the results of the ridge-regressed data are significant for every participant, and further that the improvements in performance for each word vector representation are significant (p’s < 0.05).

## 7 Conclusion

In this paper, we originally asked: (1) whether existing word vector representations used in NLP have any basis in the brain’s representational structure of lexical semantics, and, (2) if we can use features from the brain to increase the performance of existing word vector representations. Using representational similarity analysis, we found that the representational spaces of word embedding models are significantly related to the representational spaces of the fMRI data; however, a great proportion of the fMRI data’s variation beyond noise still remains uncaptured by the word embeddings. For the second question, we demonstrated that features from brain imaging data can be used to improve the performance of existing word embedding models, and that these improvements can be generalized to words which lack brain imaging data.

We introduced a novel semantic judgement acquisition technique for evaluating word embeddings in NLP: the multi-arrangement method. To measure a participant’s representational space for a set of stimuli, the method relies on the redundant distance information present in a participant’s successive arrangements of stimulus subsets. The multi-arrangement method overcomes many of the shortcomings present in existing techniques for collecting semantic judgements in NLP, namely: conflating between different types of semantic relations, relying on heuristic associations, and biasing from word order and context.

An inherent limitation of this work is the certainty of noise in fMRI data and non-invasive brain imaging techniques as a whole. We must additionally acknowledge the practicality of our results. Since we cannot expect to compile a large enough fMRI dataset of a language’s most commonly-used words, future work should explore how well our results can generalize to a larger proportion of the lexicon.

While this work is limited to concrete nouns (due to the availability of words in the fMRI dataset), future work could determine if our findings hold for other lexical items such as abstract nouns, adjectives, and verbs. Our findings may also inform the development of models aiming to learn a mapping between brain imaging data and naturalistic text stimuli, specifically, entire sentences and phrases (Sun et al., 2019; Schwartz et al., 2019; Djokic et al., 2020). Despite the most successful such attempts having used intracranial recordings (Makin et al., 2020), our results suggest that further semantic decoding progress remains to be made with non-invasive brain recordings.

Moreover, we use RSA to compare brain imaging data with computational models, and it is worth investigating applications of RSA beyond brain imaging data. For example, recent work has used RSA to compare representational spaces across computational language models and their individual components (Gauthier and Levy, 2019; Abnar et al., 2019; Chrupała and Alishahi, 2019). We also feel that the multi-arrangement method is underutilized in NLP for the shortcomings it addresses in existing semantic judgement acquisition techniques. Future research can employ the method when collecting similarity scores for a variety of textual stimuli (e.g. words, phrases).

As a whole, our work serves to motivate more research that borrows insights from neuroscience to build more appropriate computational language models.

## Footnotes

1 http://www.cs.cmu.edu/afs/cs/project/theo-73/www/science2008/data.html

2 Voxels are the three-dimensional units of fMRI data. In this dataset they are defined with a spatial resolution of 3×3×6 mm^3^ (each containing approximately one million neurons).

3 Under this threshold, the number of selected voxels ranged between 445 and 3,614 in participants. This range demonstrates the large variation in reliability across the participants’ data, which is to be expected from non-invasive brain imaging techniques.

4 Unlike in Mitchell et al. (2008) where voxel selection was done independently for each left-out fold during crossvalidation.

5 https://code.google.com/archive/p/word2vec

6 https://nlp.stanford.edu/projects/glove

7 https://fasttext.cc/docs/en/english-vectors.html

8 A rank-correlation is used to compare the RDMs because we cannot assume a linear relationship between the brain imaging data and the word vectors.

9 Noise ceiling computations were performed after rank-transforming the RDM of each participant.

10 http://www.mrc-cbu.cam.ac.uk/methods-and-resources/toolboxes/

11 http://meadows-research.com

12 We determined that 10 semantic judgements per word pair would be sufficient for controlling variability across participants. This determination was based on previous work employing the multi-arrangement method for visual stimuli, and existing word similarity/relatedness datasets.

13 The hyperparameters for each ridge regression model were optimized accordingly during training.

14 As a non-parametric test, the Wilcoxon signed-rank test made no assumptions regarding the distribution of the correlation values we provided. The resulting significance values were corrected using the Benjamini-Hochberg method.

